# Identification of specific metabolic capacities associated with major extraintestinal pathogenic *Escherichia coli* lineages

**DOI:** 10.1101/2025.09.14.675865

**Authors:** Guilhem Royer, Françoise Chau, David Vallenet, Erick Denamur, Colibafi/Septicoli/Colocoli & Coliville groups

**Affiliations:** Unité de Bactériologie, Département de Prévention, Diagnostic et Traitement des Infections, AP-HP, Hôpital Henri Mondor, Créteil 94000, France; Université Paris Cité, INSERM, IAME, Paris 75018, France; INSERM U955, Institut Mondor de Recherche Biomédicale, Créteil 94010, France; LABGeM, Génomique Métabolique, Genoscope, Institut François Jacob, CEA, CNRS, Université Evry, Université Paris-Saclay, 91057 Evry, France; AP-HP, Laboratoire de Génétique Moléculaire, Hôpital Bichat, Paris 75018, France

## Abstract

Bacterial niche colonization relies on multiple factors, among which the metabolic capacity to utilize specific substrates is pivotal, especially within complex microbiota where nutrient competition is intense. *Escherichia coli*, a gut commensal of humans and vertebrates, is also an opportunistic pathogen responsible for both intestinal and extraintestinal infections, including bloodstream infections (BSI). To identify metabolic specificities of pathogenic strains that could explain the emergence and success of major clones, we leveraged a well-characterized dataset of 1,498 genomes from both commensal and extraintestinal pathogenic epidemiologically relevant French isolates. Using a pangenomic framework, we conducted a large-scale analysis of metabolic pathways. Although metabolism was far more conserved than gene content, we found substantial metabolic diversity across the species, with over 50% of pathways being variable, mainly involving biosynthetic and degradation processes. Phylogeny had a major impact on metabolic profiles, while no specific pathway was associated with lifestyle (commensal vs. pathogenic) or BSI portal of entry. However, we identified clone-specific metabolic capacities. For example, pathways involved in 5’-deoxynucleoside recycling were enriched in the major extraintestinal pathogen (ExPEC) clone STc69. In phylogroup B2, several clone-specific pathways enabled degradation of plant-derived compounds. Notably, detailed analysis of the D-apiose degradation revealed a functional pathway strongly associated with the STc131 and STc14 pandemic clones. Overall, our study highlights multiple lineage-specific metabolic capacities that may contribute to the ecological success and dissemination of most prevalent ExPEC clones.

**Importance:** According to the nutrient-niche hypothesis, both commensal and pathogenic *E. coli* strains must exploit distinct substrates to grow and persist in their primary habitat alongside resident microbiota. Such niche differentiation also occurs between *E. coli* populations competing within the same environment and partly explains the stable coexistence of strains in the gut. With this in mind, we search for specific associations between metabolic pathways and strain origin. However, metabolic profiles were predominantly shaped by phylogeny, reflecting the species’ clonal structure and the close link between phylogenetic background and lifestyle. Among the lineage-specific determinants, we identified several pathways associated with worldwide spread clones responsible for bloodstream infections, supporting the existence of clone-specific strategies for niche adaptation.

## Introduction

Metabolism is key to bacterial adaptation to nutritional environments. This has been extensively studied in *Escherichia coli*, a species that includes both commensal, intestinal (InPEC) and extra-intestinal pathogenic strains (ExPEC) (1, 2). The primary habitat of *E. coli* is the intestinal tract of mammals and many other vertebrates (2, 3). Within this reservoir, the nutrient-niche hypothesis proposes that *E. coli* strains can coexist with resident microbiota by exploiting distinct nutrients (4). According to this theory, *n* populations may coexist if *n* distinct substrates are available, with population sizes controlled by substrate concentrations. However, not only nutrient availability but also strain-specific affinity for given nutrients contributes to colonization success and coexistence (5, 6). Spatial variation within the gut may add a third layer of complexity, whereby local microenvironments influence strain distribution (6, 7). Importantly, the nutrient-niche hypothesis extends beyond the intestinal tract and applies to other environments encountered during infection. For example, the survival and growth of uropathogenic *E. coli* (UPEC) in urine is facilitated by the *dsdCXA* locus, which enables D-serine utilization, an amino acid abundant in urine (8, 9).

*E. coli* exhibits a clonal population structure with eight major phylogroups, namely A, B1, B2, C, D, E, F, and G (1, 2, 10), as well as cryptic clades that are phenotypically indistinguishable but genetically divergent (11). This structure is supported by both genetic distances (12, 13) and gene content patterns (14). Consequently, *E. coli* harbors a pan- and core-genome not only at the species level but also within phylogroups, revealing a fractal-like genetic organization. Within phylogroups, subpopulations are commonly delineated using multilocus sequence typing (MLST) (15), sometimes further grouped into biologically meaningful sequence type complexes (STcs).

Phylogroups are non-randomly associated with the different *E. coli* hosts, and host diet may shape their distribution (3), supporting the relevance of phylogroup-specific nutrient-niche adaptation. Similarly, some ST/STcs are strongly associated with specific hosts, such as STc1l7 (phylogroup G) with poultry (16), and STEC O157:H7 ST11 (phylogroup E) with livestock reservoirs (17, 18). Longitudinal studies in humans (19, 20) also revealed phylogroup-dependent colonization patterns. These findings support, in addition to niche differentiation, the existence of a trade-off between colonization and residence: phylogroups B2 and F tend to be long-term residents of the gut, while A, B1, and D display strong colonization abilities.

Another key aspect of the *E. coli* population is the strong link between phylogroups and lifestyles: commensals are mainly in phylogroups A and B1, while ExPEC strains are largely found in B2, D, and F (1), these associations being not mutually exclusive. This overlap, potentially resulting from epistatic interactions (1), makes it difficult to disentangle phylogroup from lifestyle-associated traits, such as metabolic pathways. Furthermore, identifying metabolic features specific to lifestyle is challenging because commensals, InPEC and ExPEC all inhabit the gut and must compete for similar nutrients. These difficulties along with the major footprint of phylogeny on metabolism were already evident in a pioneering study of pan and core-metabolism of 29 *E. coli* strains, including *Shigella* strains (21), which found that metabolic profiles were more strongly associated with phylogroups than with pathotypes. Only a limited set of reactions were preferentially associated with either commensals or ExPEC strains, and none were entirely specific.

Subsequent studies also aimed to identify phylogroup- or pathotype-specific metabolic profiles. For instance, using genome-scale metabolic reconstructions of 55 *E. coli* strains, Monk *et al*. identified a few pathways that could differentiate ExPEC from commensals (22). However, presence/absence patterns of many of them were also specific of phylogroup B2, consistent with the distinct gene repertoire of this group (23). Notably, the absence of pathways such as fructoselysine and psicoselysine (Amadori products) degradation, as well as 3-(3-hydroxyphenyl)propanoate catabolism was mainly observed in ExPEC, but it is also a hallmark of B2 strains (23, 24). Yet, exceptions exist. For instance, the pandemic ExPEC ST131 and the atypical B2 commensal ST452 do carry the latter pathway (19, 25), highlighting intra-phylogroup heterogeneity. Indeed, as expected due to phylogroup-specific pan- and core-genomes (14), finer-scale analyses also revealed ST-specific metabolic profiles (26, 27) adding another layer of complexity to attempts to link metabolism with lifestyle.

To date, knowledge of *E. coli* population metabolic specificities is often derived either from high-quality but limited datasets (21, 22, 28) or from larger, more comprehensive but heterogeneous collections (25). Notably, the absence of metadata regarding strain origin and context of isolation may often limit the epidemiological relevance and interpretability of findings. In this context, we analyzed a large and well-defined genome collection of *E. coli* strains, including both commensals and ExPEC, gathered during prospective, multicentric studies in France. Using a pangenome approach coupled with metabolic pathway predictions, we explored metabolic features associated with both lifestyle and phylogenetic background, with particular focus on the dominant STcs (i.e. STc131, 95, 73, 69, 10 and 14) causing bloodstream infections (BSI). Our aim was to identify metabolic traits that may be associated with strain sources and/or contribute to the epidemiological success of some STc, under the hypothesis that metabolism is a key determinant of ExPEC major clone’s emergence and infection potential.

## Methods

### Commensal and pathogenic strain genome datasets

We analysed *E. coli* genomes from both commensal (n=370) and extraintestinal pathogenic (ExPEC) (n=1128) human strains, all isolated in France across a time period ranging from 2000 to 2017. Commensal strains (29) consisted of five collections obtained from feces or rectal swabs and gathered from 2000 to 2017: ROAR in 2000 (n = 50) (30) and LBC in 2001 (n = 27) (31) in Brittany; PAR in 2002 (n = 27) (31), Coliville in 2010 (n = 246) (32) and CEREMI in 2017 (n = 20) (33) in the Paris area. Pathogenic strains were obtained from three collections: Colibafi (n=367) (34) and Septicoli (n=545) (35) which correspond to strain isolated from BSI in the Paris area in 2005 and 2016-17, respectively; Colocoli, a collection of *E. coli* pulmonary strains (n=216) from mechanically ventilated patients in French intensive care units (36). Among pathogenic strains, the source of infection was urinary (n=470), digestive (n=281), pulmonary (n=235) or other (n=142) [unknown (n=57), multiple (n=39), catheter (n=29), skin (n=9), gynecologic (n=4), surgical site (n=4)]. Phylogroups, Multi Locus Sequence Type (MLST), ST complex (STc) were retrieved from the corresponding studies. All genomes were short-read sequenced on Illumina plateforms. In sum, these strains can be considered as epidemiologically relevant. Detailed information, including the corresponding bioprojects, are available in Table S1.

Independence between phylogroups and sources was tested using a chi-square test. As a post-hoc test, adjusted Pearsons residuals were analysed after Bonferroni multiple-test correction.

### Pangenome construction

Genomes were annotated with Pyrodigal v3.2.1 (37), Aragorn v1.2.41 (38) and Infernal v1.1.4 (39) and the pangenome was built using PPanGGOLiN v2.0.0 (40), applying thresholds of 80% amino acid identity and alignment coverage to define protein families. The pangenome file is available on zenodo (https://doi.org/10.5281/zenodo.17104470). From the multiple sequence alignment of core genes (option “MSA” of PPanGGOLiN), a phylogenetic tree was computed with iqtree v1.6.12 (41) and the GTR+F+I+G4 model as previously described (23). Patristic distances between all genome pairs were determined from this tree with the function “cophenetic” from R package “ape” (42).

### Panreactome and pathway prediction

To determine Gene-Protein-Reaction (GPR) associations at the pangenome level, we applied a three-step approach in which representative protein sequences from each pangenome family were aligned against reference protein sequences using Diamond v2.1.8 (43) in ‘ultrasensitive’ mode, with MetaCyc v27.0 (44) serving as the reference database for reactions and pathways. First, representative pangenome sequences were compared with *E. coli* K-12 protein sequences from EcoCyc v27.0 database (45). Only matches with at least 80% identity and coverage were considered, and EcoCyc reaction annotations with equivalents in MetaCyc were transferred to the corresponding pangenome families. Second, for representative proteins without K-12-related reactions, we compared them against MetaCyc protein sequences to transfer corresponding reaction annotation, considering only best hits with a minimum identity of 40% and coverage of 80%. Finally, for the remaining proteins, we ran kofamScan v1.3.0 (46) with KofamKoala database (version 2023-10-02), retaining only hits above the score threshold defined for each KEGG Orthology (KO) group. To annotate pangenome families, KEGG reactions associated with each KO group were mapped to the corresponding MetaCyc reactions. If no cross-reference was found, only EC (Enzyme Commision) numbers of KO were used to assign enzymatic activities.

The resulting panreactome, comprising GPR associations described at the pangenome family level with either MetaCyc reactions or EC numbers, was used as input of Pathway Tools v27.0 (47) which was run by command line with default parameters and using PathoLogic file format for annotations (https://doi.org/10.5281/zenodo.17107335). For each predicted pathway, a completion value at the pangenome level was computed by dividing the number of predicted reactions by the total number of reactions in the pathway, excluding spontaneous reactions and also orphan ones (i.e. reactions not associated with any known gene in the MetaCyc database). Pathway completions were also computed for individual genomes from the pangenome gene family presence/absence matrix. A pathway was considered present in a given organism when its completion reached more than 50% of the maximum completion value observed at the pangenome level.

### Metabolic distances and clustering

Metabolic distances were computed as Manhattan distances between genomes based on presence/absence of the predicted pathways. A linear regression using “lm” from R package “stats” was used to search for a correlation between metabolic and patristic distances.

We also performed a hierarchical clustering of genomes based on metabolic distances, considering only pathways with frequencies ranging from 5 to 95%, with the Ward D2 method using “hclust” from R package “stats”.

### Multifactorial correspondence analysis

A factorial multiple correspondence analysis (MCA) was performed with FactoMineR (48) using the presence/absence of each pathway as active variables. Only pathways present in 5% to 95% of genomes were considered. Strain origins and phylogroups were used as illustrative variables. The results were plotted using Factoextra (49), considering the first two eigenvalues. The same analysis was run using the presence/absence of reactions as active variables.

### Genomic characterization of D-apiose degradation gene cluster

To characterize D-apiose degradation gene cluster, we used the MicroScope platform (50) and Clinker (51) to compare gene clusters from *Pectobacterium carotovorum* WPP14 (Refseq accession number: GCF_013488025.1), where the pathway has been previously described (52), *Escherichia fergusonii* ATCC35469 (Refseq accession number : GCF_000026225.1), *E. coli* H1-004-0008-M-Y (ST131 025b-H4-fimH30 - Clade C1), *E. coli* H1-003-0083-B-J (ST131 025b-H4-fimH30 - clade C2) and *E. coli* H1-002-0016-H-R [ST1193 (STc14)]. This approach enables the comparison of each coding sequence at the protein level and the analysis of synteny conservation.

In a second step, we analysed the location of the gene cluster among all *E. coli* complete genomes available in RefSeq on September 19, 2022 (n=2302) (53), as well as other *Escherichia* non-coli species (n=167). Nucleic sequences spanning from *entH* to *cusS* were extracted and annotated using Bakta v1.9.1 (54). From the pangenome computed with PPanGGOLiN, a subgraph of this genomic region was extracted and visualized with Gephi (55). The pangenome file is available on zenodo (https://doi.org/10.5281/zenodo.17107020). Using this complete genome dataset, we also conducted three phylogenetic analyses with iqtree: one based on the species core gene alignment, one on the whole *entH-cusS* region, and one on the apiose gene cluster. The core gene phylogeny was inferred using the GTR+F+I+G4 model, while the most appropriate model for the other two alignments was selected using ModelFinder (56).

### Large-scale screening of D-apiose degradation gene cluster

We performed a large-scale search for the presence of the 10 genes of the apiose cluster among the 2,440,377 genomes assemblies from the database AllThebacteria (57). We built an *ad-hoc* database for abricate (58) using nucleic sequences of the genes from *E. coli* SE15 (B2, ST131) (RefSeq accession number: NC_013654.1; gene locus tag ECSF_RS02670 to ECSF_RS02715). Then, we screened genomes using minimal nucleotidic identity and alignment coverage of 50%. The pathway was considered present when at least one gene homolog was detected in the compared genomes.

### Growth under D-apiose-containing M9 minimal media

*P. carotovorum* WPP14, *E. fergusonii* ATCC 35469, three *E. coli* strains carrying the apiose gene cluster (H1-004-0008-M-Y, H1-003-0083-B-J and H1-002-0016-H-R), and *E. coli* K-12 MG1655 were grown aerobically in LB medium at 37⍰ °C overnight. Bacterial cells were then washed and diluted 1:1,000 into M9 minimal medium supplemented with 10⍰mM apiose (Omicron). Growth was monitored over 48 hours at 37⍰°C under continuous shaking by automatic measurement of optical density (OD) at 600⍰nm.

## Results

### An epidemiologically relevant collection with lifestyle-specific population genetic structure

The *E. coli* collection used in this study comprises 1,498 genomes and is representative of the diversity of commensal and ExPEC strains isolated in humans living in high income countries (1, 2). A greater diversity is observed in commensal strains, whereas ExPEC strains are pauciclonal (Table S2), particularly for urinary isolates where the ten most prevalent STc represent more than 80% of the strains. Overall, the collection covers the eight main *E. coli* phylogroups, along with one strain from phylogroup H and eight from *Escherichia* clades (Table S3). Several previously reported associations between phylogroups and isolation sources were observed (2, 59–62). Phylogroups A and B1 were significantly underrepresented among urinary isolates, and phylogroups A, E and F more frequently associated with commensal strains (Table S3, Table S4). In contrast, phylogroups B2 were significantly less common among commensal isolates, but enriched in urinary isolates.

### A collection with a high genomic and metabolic diversity

Assuming that we have an epidemiologically valid dataset, we analyzed the diversity of the whole collection in terms of gene families, reactions, and metabolic pathways (Figure 1). The rarefaction curves revealed relatively closed panreactome (Figure S1A) and panmetabolic pathway repertoires (Figure 1A), comprising 3,023 reactions and 487 predicted pathways, in contrast to a much larger and more open pangenome consisting of 37,236 gene families. Reactions and pathways appeared more conserved than genes, reaching core values of 1,360 reactions (44.99%) and 243 pathways (44.90%), respectively, compared with 1,400 core genes (3.76%) (Figures 1B and S1B). The frequency distribution of reactions and pathways displayed a U-shape (Figures 1C and S1C), inverted compared to the typical gene frequency distribution (63), with a higher proportion of conserved elements compared to unique or rare ones. A large number of variable pathways were associated with the “Biosynthesis” and “Glycan Pathways” categories, with an overrepresentation of pathways involved in O-antigen biosynthesis, present in 27/60 (45.0%) and 27/29 (93.1%) of the variable pathways in these categories, respectively (Figure 1D). Another major subset of the variable pathways (62/161, 38.5%) was linked to “Degradation” processes.

**Figure 1.**
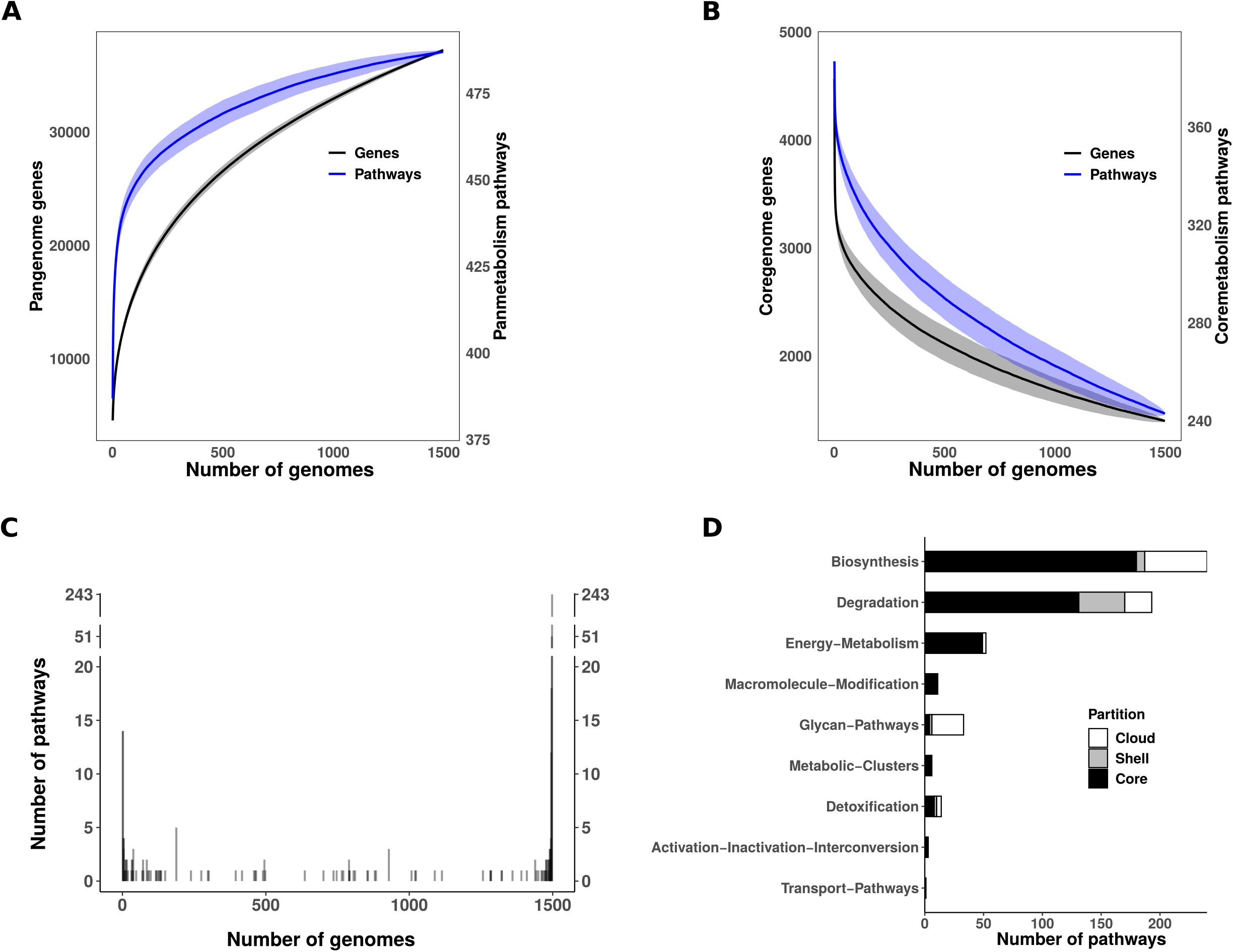
*E. coli* core- and pan-metabolism. Evolution of (A) the pangenome and panmetabolic pathways, and (B) the coregenome and coremetabolic pathways as a function of the number of included genomes. To account for genome variability, 1,000 random permutations were performed at each step of genome addition. The resulting mean number of genes and pathways are shown in black and blue, respectively. Shaded areas represent the standard deviation. (C) Frequency of pathways across the 1,498 genomes analyzed. Pathways on the left side of the graph are present in only one genome (n=14, 2.87% of panmetabolic pathways), while those on the right side are found in all genomes (n=243, 49,90% of panmetabolic pathways). For the sake of readability, the y-axis is broken. (D) Distribution of pathways among the main metabolic functional categories defined by MetaCyc. Bar plots are colored according to pathway frequency. Core (black), shell (grey) and cloud (white) partitions correspond to pathways with frequencies f ≥ 95%, 15% ≥ f > 95% and f <15%, respectively.

### Metabolic diversity is mainly driven by phylogeny

In a second step, we compared the sizes of the pangenome, panreactome, and panmetabolic pathways relative to the number of strains, taking into account their origin and phylogroup. Overall, when all sources were included, no significant correlation was observed between the number of genomes and the number of genes, reactions, or pathways (Figures 2A, S2A and S2B). However, excluding strains of urinary origin markedly improved the correlation, which became significant for both reactions and pathways (Figure 2B, S2C and S2D). When analyzed by phylogroups, significant correlations were observed for gene, reaction, and pathway counts (Figures 2C, S2E and S2F). Notably, excluding phylogroup A strains further strengthened these correlations in most cases (Figures 2D, S2G and S2H). Together, these results suggest a reduced metabolic and genomic diversity among strains isolated from urinary-source bacteremia, while strains from phylogroup A appear to harbor increased diversity.

**Figure 2.**
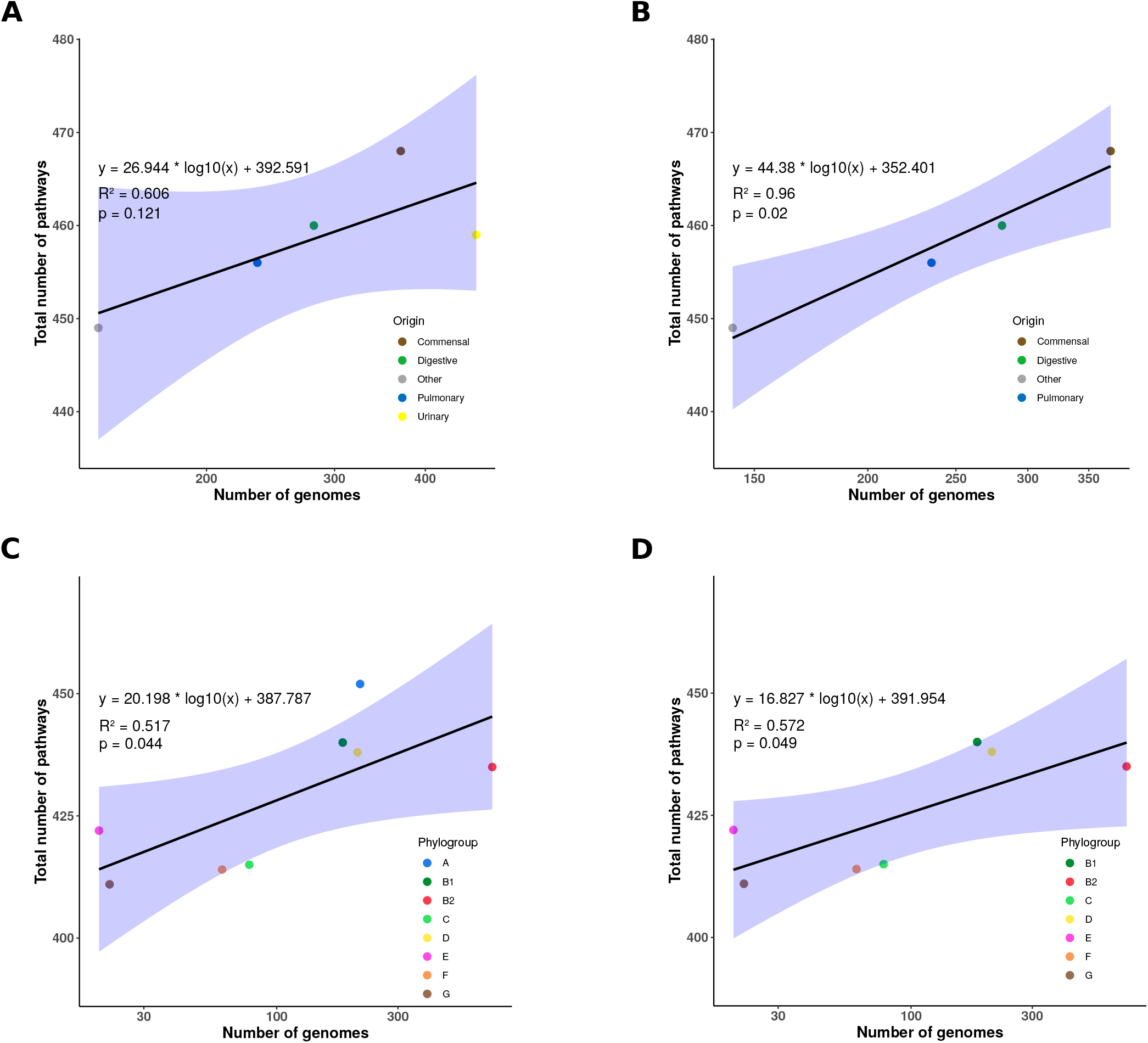
Correlation between the number of pathways and genomes. Total number of pathways as a function of the number of genomes: (A) from all sources, (B) excluding those of urinary origin, (C) from all phylogroups, and (D) excluding phylogroup A. Data points are colored according to their origin or phylogroup. The regression line is shown in black, with the 95 % confidence interval in blue. Each graph includes the linear regression equation, the coefficient of determination (R^2^) and the p-value.

We then investigated whether specific pathway and reaction presence/absence profiles could segregate strains based on their phylogroup and/or isolation source. To this end, we performed a multifactorial correspondence analysis (MCA) using the presence/absence matrix of metabolic pathways (Figure 3A) or reactions (Figure S3) across the 1,498 strains. The first two dimensions of the MCA captured a substantial proportion of the overall variability, accounting for 42.6% for pathways and 41.8% for reactions. When illustrative variables were mapped onto the MCA plots, metabolic diversity appeared primarily structured by phylogeny rather than origin. Specifically, the first dimension mainly separated phylogroup B2 from phylogroups A, B1 and C, while the second dimension distinguished phylogroups D, E and F from the others. No clear separation was observed between strains based on their origins along these axes. These findings are consistent with the observed correlation between metabolic and patristic distances computed across all genome pairs (Figure 3B), excluding clades. The inclusion of *Escherichia* clades disrupted the correlation, likely due to their substantial genetic divergence from *E. coli sensu stricto* (Figure S4).

**Figure 3.**
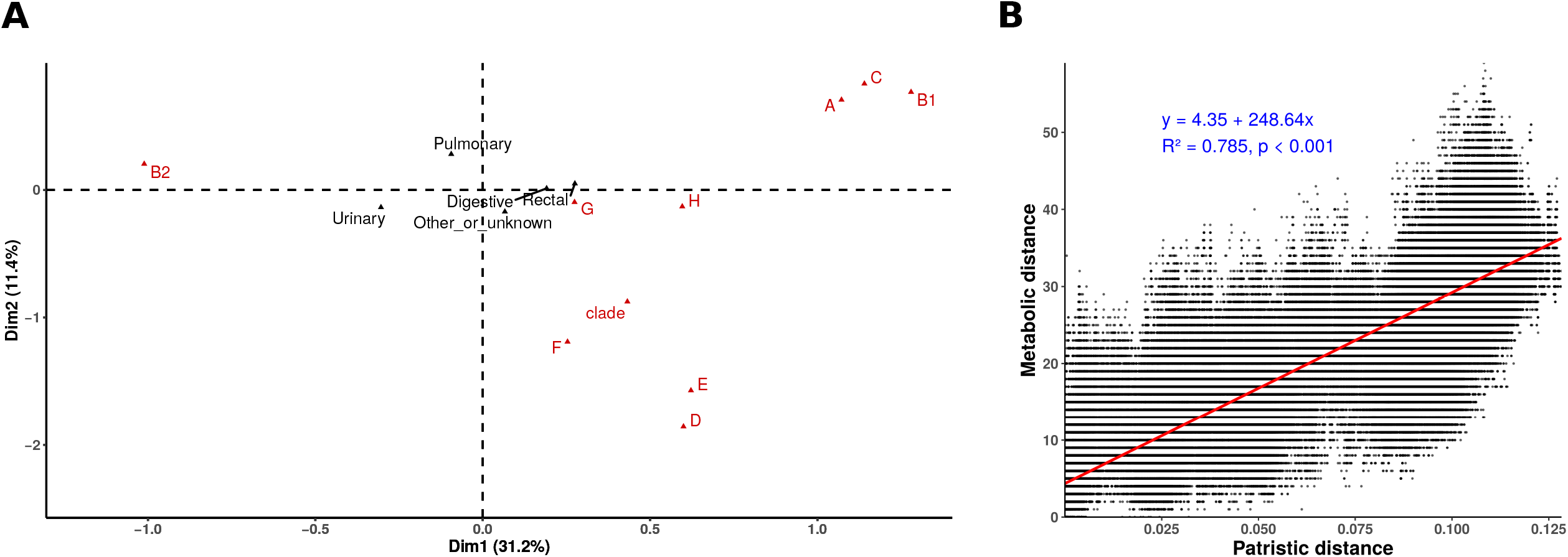
Correlation between phylogeny, origin and metabolism. (A) Multiple correspondence analysis of pathway occurrences among the 1,498 genomes analysed. The x- and y-axes represent the first dimensions, which together account for 42.6% of the variability. Phylogroups (in red) and origin (in black) are shown as illustrative variables. (B) Correlation between patristic and metabolic distances. Patristic distances represent the branch lengths between genome pairs in the coregene-based phylogenetic tree. Metabolic distances are Manhattan distances computed from the binary pathway presence/absence matrix. Each point represents a pair of genomes. The regression line is shown in red, while the linear regression equation, coefficient of determination (R^2^) and p-value are shown in blue. Only strains belonging to *E. coli* sensu stricto are included in panel B (i.e. *Escherichia* clade strains are excluded).

### A limited number of pathways separate phylogroups and major STcs responsible for BSI

The lack of a global correlation between isolation source and pathway presence/absence does not rule out the possibility that a small subset of metabolic pathways play specific roles in adaptation or pathogenesis. Indeed, analyzing all pathways collectively may have masked more subtle associations. To explore this, we focused on metabolic pathways with intermediate frequencies (5–95%), excluding both highly conserved and very rare ones. Analysis of the presence/absence profiles of these 64 pathways did not reveal any segregation based on strain origin (Figure 4). Instead, the patterns strongly mirrored phylogroup classification. Phylogroup B2 appeared particularly distinct from others, largely due to the absence of multiple pathways.

**Figure 4.**
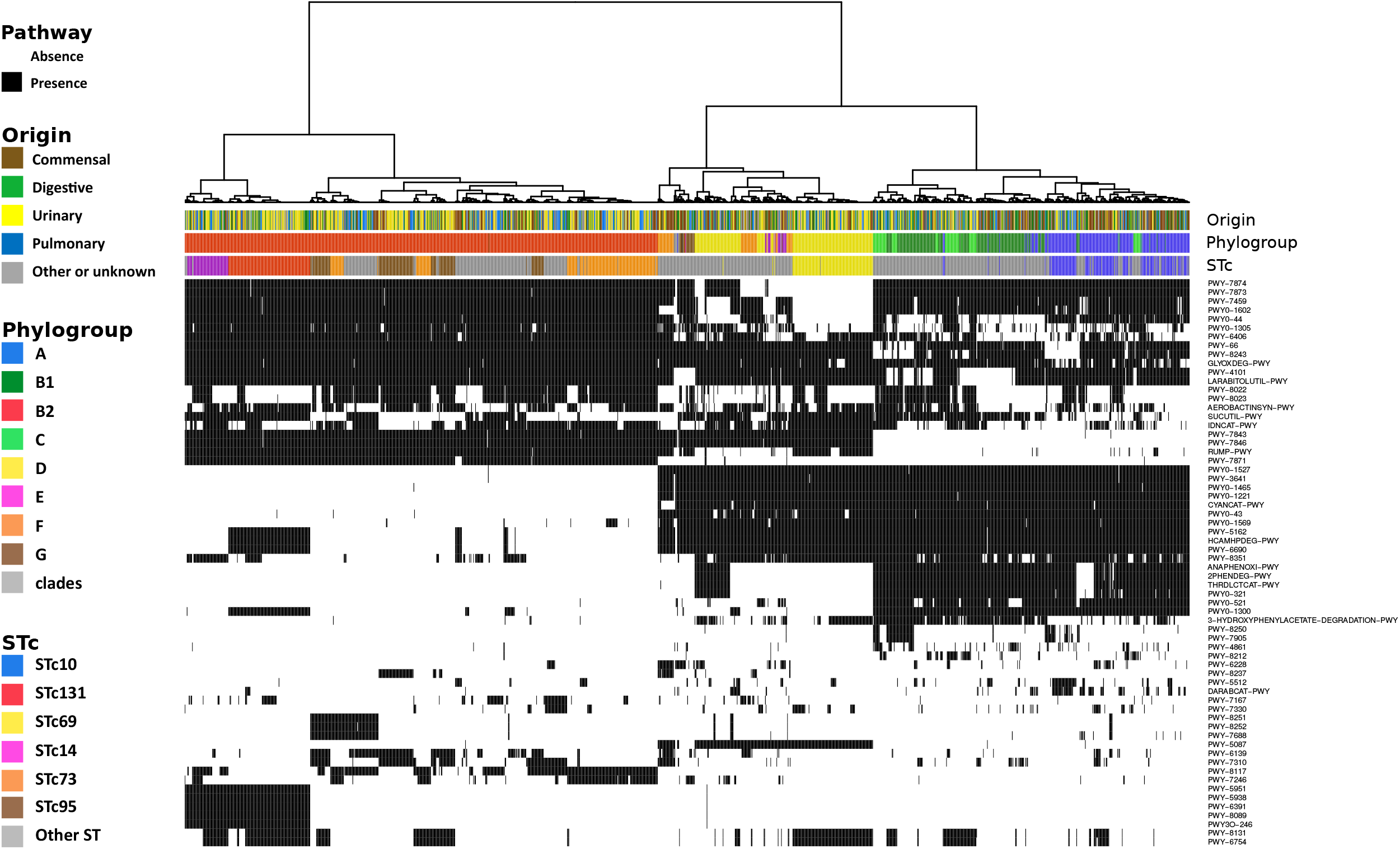
Hierarchical clustering of metabolic pathways and associations with origins, phylogroups and major ExPEC STcs. The dendrogram above the heatmap represents the clustering of genomes according to their pathway content. Clustering was performed with the Ward D2 method on Manhattan distances based on the presence and absence of metabolic pathways having a frequency between 5% and 95%, shown in black and white on the heatmap, respectively. From top to bottom, the color strips indicate the origin, phylogroup and STc of each strain. Only the most prevalent STc associated with bacteremia (76), along with the emerging high-risk clone STc14 (78), are highlighted in color; all other STs are shown in grey.

At a finer resolution, specific metabolic signatures were observed in major STcs associated with BSI. Notably, STc131 and STc14 (both from phylogroup B2), as well as STc69 (from phylogroup D), formed well-defined clusters on the heatmap, suggesting conserved and distinctive metabolic profiles within these clinically significant lineages.

To identify a limited set, or ideally unique, metabolic pathways strongly associated with pathogenicity, BSI portal of entry, or phylogroup, we next examined the frequency of individual metabolic pathways across these variables. We specifically searched for pathways that were overrepresented (≥80% of frequency) in at least one group and underrepresented (<20% of frequency) in at least another, across the following comparisons: (i) commensal versus pathogenic strains, (ii) strains stratified by BSI portal of entry, and (iii) phylogroups. No pathway fulfilled these criteria when comparing lifestyles (commensal, pathogen or BSI portal of entry). However, 34 pathways were found to be either overrepresented or underrepresented in at least one phylogroup (Table S5). All but one of these pathways were associated with degradation processes. Interestingly, some of these pathways, such as heme degradation IV and V (PWY-7843, PWY-7846), encoded by the *chu* gene cluster, are the basis of Clermont’s phylogrouping method (64, 65) and separate A, B1, C (absence) from others (presence).

We used the same approach to investigate whether specific pathways were enriched within particular STcs relative to the rest of their respective phylogroup. No such pathways were found for STc10 compared to other phylogroup A strains, in line with the extensive genetic and metabolic diversity of this group. In contrast, three pathways were found to be specifically enriched in STc69 (phylogroup D) compared to other strains within the same phylogroup: S-methyl-5’-thioadenosine degradation I, 5’-deoxyadenosine degradation II, and formaldehyde oxidation I (Table S6). The first two have been recently described in a dihydroxyacetone phosphate shunt, enabling the recycling of 5⍰-deoxynucleosides which may accumulate in urine and blood in humans (66, 67).

Also, among the major STcs responsible for extraintestinal diseases within phylogroup B2, namely STc131, STc73, STc95, and STc14, we identified 13 pathways with distinct presence/absence patterns (Table 2). Most of these were involved in degradation (8/13), followed by biosynthesis processes (3/13). Three of the STc131-specific pathways were linked to the *mhp* gene cluster (HCAMHPDEG-PWY, PWY-5162, PWY-6690) (19). STc95 was characterized by the presence of the CMP-N-acetylneuraminate biosynthesis II pathway, consistent with the known high frequency of type II capsule production in this lineage (68). Other STc-specific pathways included 2-O-α-mannosyl-D-glycerate degradation (STc131), pectin degradation II (STc73), D-glucosaminate degradation (STc95), and sulfoquinovosyl diacylglycerides and sulfoquinovosyl glycerol degradation (STc14). Additionally, five pathways were shared between STc131 and STc14, with identical frequencies. Closer inspection revealed that these were all predicted based on reactions involved in D-apiose degradation, suggesting a shared acquisition of this specific catabolic capacity.

**Table 1.** Number of core and variable genes, reactions and pathways according the origin and phylogeny of strains

|  |  | Genes |  |  | Reactions |  |  | Pathways |  |  |
| --- | --- | --- | --- | --- | --- | --- | --- | --- | --- | --- |
|  | Number of genomes | Core* | Shell** | Cloud*** | Core* | Shell** | Cloud*** | Core* | Shell** | Cloud*** |
| All | 1498 | 3260 | 2373 | 31603 | 2145 | 376 | 502 | 360 | 46 | 81 |
| Origin |  |  |  |  |  |  |  |  |  |  |
| Commensal | 370 | 3213 | 2317 | 21760 | 2139 | 367 | 428 | 358 | 48 | 62 |
| Pathogenic | 1128 | 3276 | 2380 | 25702 | 2144 | 382 | 429 | 360 | 51 | 65 |
| Urinary | 470 | 3316 | 2458 | 16076 | 2151 | 377 | 319 | 361 | 50 | 48 |
| Digestive | 281 | 3254 | 2284 | 17066 | 2139 | 371 | 366 | 360 | 44 | 56 |
| Pulmonary | 235 | 3292 | 2219 | 12672 | 2145 | 368 | 313 | 361 | 43 | 52 |
| Other | 142 | 3273 | 2385 | 11660 | 2144 | 385 | 261 | 360 | 49 | 40 |
| Phylogeny |  |  |  |  |  |  |  |  |  |  |
| A | 213 | 3413 | 1575 | 16868 | 2204 | 200 | 385 | 369 | 31 | 52 |
| B1 | 182 | 3610 | 1357 | 13465 | 2259 | 160 | 292 | 380 | 20 | 40 |
| B2 | 705 | 3653 | 1900 | 13651 | 2250 | 192 | 263 | 376 | 22 | 37 |
| C | 78 | 3747 | 1364 | 4782 | 2276 | 154 | 118 | 381 | 21 | 13 |
| D | 208 | 3617 | 1653 | 9811 | 2259 | 194 | 241 | 376 | 25 | 37 |
| E | 20 | 3658 | 2247 | 3664 | 2250 | 162 | 152 | 375 | 20 | 27 |
| F | 61 | 3559 | 2052 | 4367 | 2246 | 181 | 119 | 371 | 28 | 15 |
| G | 22 | 3778 | 1358 | 2880 | 2293 | 87 | 112 | 382 | 10 | 19 |
| H | 1 | 4483 | NA | NA | 2315 | NA | NA | 382 | NA | NA |
| Clades | 8 | 3181 | 1921 | 1952 | 2066 | 330 | 59 | 345 | 46 | 7 |

\*Core corresponds to elements present in at least 95% of the genomes \*\*Shell corresponds to elements present in at less than 95% and at least 15% of the genomes \*\*\*Cloud corresponds to elements present in less than 15% of the genomes

**Table 2.**
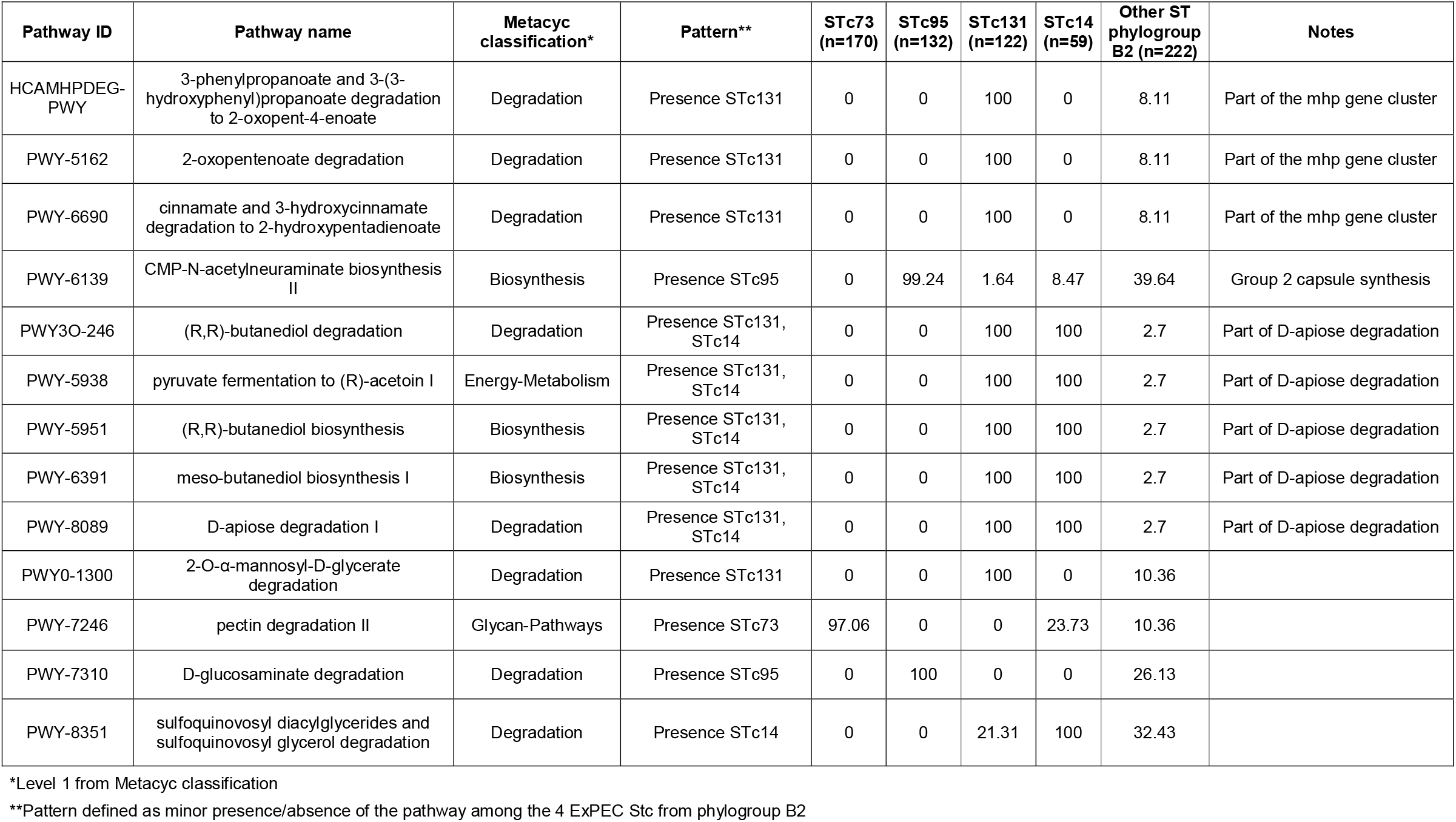
Pathways specifically absent/present among the major ExPEC Stc from phylogroup B2. Frequencies (%) of pathways are presented for each major ExPEC Stc and other ST from phylogroup B2.

In sum, only a few metabolic pathways appear to be specifically associated with the major STcs responsible for bacteremia. Strikingly, in phylogroup B2, most of these pathways are involved in the degradation of plant-derived compounds, pointing to a possible ecological adaptation of these successful clones through the acquisition of specialized catabolic functions. Two other notable pathways identified in STc69 may contribute to growth outside the intestinal niche, particularly during the infection process, for instance, in urine.

### Characterization of the D-apiose degradation pathway in *Escherichia sp*

W*e* then focused on the D-apiose degradation pathway (Figure 5A), as it emerged as a core metabolic feature uniquely shared between the two most recent pandemic *E. coli* clones, STc131 and STc14, and absent in the rest of *E. coli* species. Screening complete genomes from the RefSeq database confirmed its presence in 100% of STc131 (n=126) and STc14 (n=22) strains, as well as in 19 of 74 (25.68%) *E. fergusonii* genomes. A few other sequence types also carried the apiose pathway gene cluster, including ST583 (n=2), ST788 (n=2), ST636 (n=1), ST7898 (n=1), and ST122 (n=1) (Figure S5). All of them belong to phylogroup B2.

**Figure 5.**
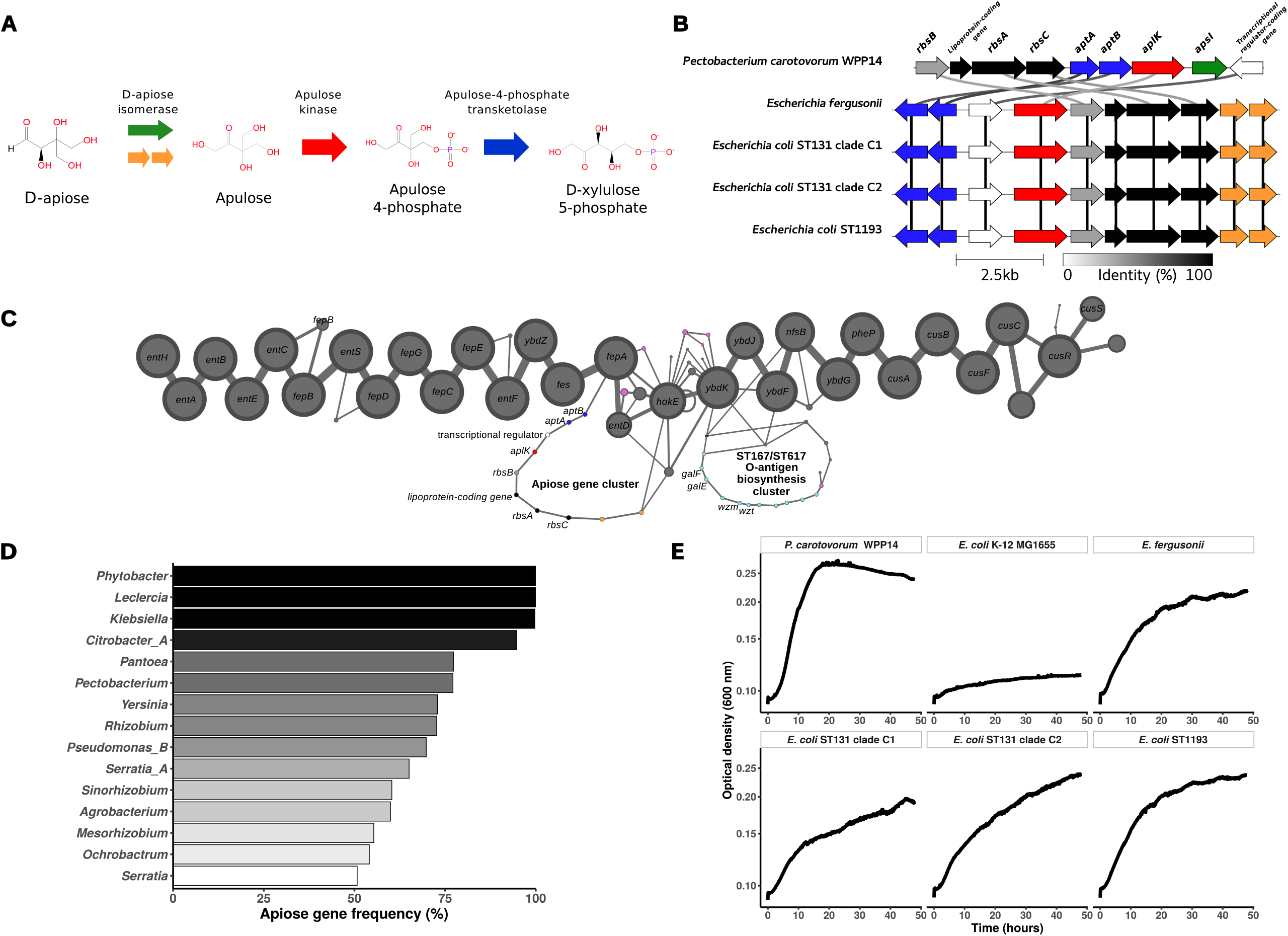
Genomic and phenotypic characterization of the D-apiose degradation pathway. (A) Schematic representation of the D-apiose degradation pathway (MetaCyc ID: PWY-8089). Reactions are indicated by colored arrows, adapted from the color scheme proposed by Carter *et al*. (69). The two putative oxidoreductases, hypothesized to functionally replace the D-apiose isomerase in Escherichia genus, are highlighted in orange. (B) Comparison of the apiose gene cluster in *Pectobacterium carotovorum* WPP14 (Assembly accession: GCF_013488025.1), *Escherichia fergusonii* ATCC 35469 (Assembly accession: GCF_000026225.1), *E. coli* H1-004-0008-M-Y (ST131 025b:H4-fimH30 – clade Cl), *E. coli* H1-003-0083-B-J (ST131 025b:H4-fimH30 – clade C2), and *E. coli* H1-002-0016-H-R (ST1193). Genes are depicted as arrows, with colors corresponding to the reactions in panel A. Shading between genes in neighboring clusters indicates protein sequence identity (0% = white; 100% = black). The figure was generated using Clinker (51). (C) Pangenome graph representation of the genomic region spanning *entH* to *cusS* in complete genomes of *E. coli* (n = 2,302) and other *Escherichia* species (n = 167). Nodes represent gene families, and edges represent their genomic neighborhood. Node and edge sizes are proportional to their frequency in the dataset. Nodes corresponding to the apiose gene cluster are colored as in panel B; transposase-related nodes are shown in pink, and those corresponding to the ST167/ST617 O-antigen biosynthesis cluster are shown in light blue. For the sake of readability, only edges found in more than 2% of genomes are represented. (D) Large-scale genome screening for the presence of apiose cluster-related genes. Bar plots show the frequency of apiose degradation-associated genes in the AIITheBacteria database. Screening was performed using abricate with a minimum identity and coverage threshold of 50%. Only genera represented by more than 50 genomes and with at least one gene with a frequency above 50% are shown. The taxonomy, retrieved from AllTheBacteria database, is determined based on GTDB (82). (E) Growth curves using D-apiose as the sole carbon source. Growth was monitored by automated OD600 measurements over 48 hours at 37⍰°C. *P. carotovorum* WPP14 and *E. coli* K-12 MG 1655 were used as positive and negative controls, respectively. All other tested strains, carrying the apiose gene cluster, correspond to those shown in panel B.

Gene cluster comparison among representative strains of *E. fergusonii* (ATCC 35469), *E. coli* STC131 clade Cl (H1-004-0008-M-Y), STc131 clade C2 (H1-003-0083-B-J), and STc14 (Hl-002-0016-H-R, ST1193) revealed an identical gene organization (Figure 5B). While similar to the cluster described in *Pectobacterium carotovorum* WPP14 (69), the *Escherichia* version did not encode the *apsI* gene coding for a D-apiose isomerase. Instead, it featured two coding sequences (CDSs) encoding putative oxidoreductases, which we hypothesize functionally replace *apsl* in this genus.

To further investigate the genomic context of the apiose cluster, we constructed a pangenome from all complete *Escherichia* genomes (Table S7). In all cases, the cluster was located downstream of the *enterobactin* biosynthesis genes (Figure 5C). Both upstream and downstream regions were highly conserved across the *Escherichia* genus. Interestingly, a distinct locus encoding an O-antigen biosynthesis cluster, previously described as specific to ST167-ST617 from phylogroup A (70), was also found in this genomic region. Phylogenetic analyses of the apiose gene cluster and its flanking regions (Figures S6 and S7) revealed strong congruence with the core-genome phylogeny (Figure S5). Moreover, within the STc131, the apiose gene cluster phylogeny perfectly matched the main clades A, B and C1/C2 (Figures S6).

To evaluate the broader distribution of apiose-related genes, we then screened the AllTheBacteria database (n=2,440,377 genome assemblies). Homologs were detected in 99 genera and 491 species, most of which are environmental or plant-associated bacteria such as *Phytobacter, Leclercia, Klebsiella, Citrobacter, Pectobacterium*, and *Rhizobium* (Figure 5D). To investigate a potential progenitor, we compared apiose-related genes from *Escherichia* and *non-Escherichia* genera. However, none of the *non-Escherichia* genera displayed genes sharing more than 90% nucleotide identity with those of *E. coli*.

Finally, to experimentally validate the predicted phenotype, we performed growth assays in M9 minimal medium supplemented with D-apiose as the sole carbon source. While *E. coli* K-12 failed to grow, the positive control *(P. carotovorum)* and all apiose cluster-positive strains, including *E. fergusonii* (ATCC 35469), STc131 clade C1 and C2, and STc14, successfully grew under these conditions, consiesnt with a functional D-apiose utilization.

Together, these data demonstrate the specific and functional presence of the D-apiose degradation pathway in STc131 and STc14. The highly conserved cluster organization across these lineages, including the distantly related *E. fergusonii*, along with its congruent phylogeny and substantial divergence with copies from *non-Escherichia* genus, support an ancient acquisition event in the evolutionary history of *Escherichia*, probably followed by multiple independent losses in other lineages.

## Discussion

*E. coli* is the main causative agent of both community- and hospital-acquired BSI (71), associated with a substantial burden (72). Despite advances in care, mortality remains high, ranging from 10 to 30% (35, 72−74). Yet, *E. coli* is also a gut commensal in humans and other vertebrates (2, 3), highlighting its high intrinsic capacity for colonization. Indeed, colonization of the digestive tract is a prerequisite for both commensal and pathogenic strains. While so-called virulence factors have been shown to increase fitness of ExPEC in the gut (75), differences in metabolic capacities may also play a pivotal role in their successful intestinal colonization. Recent longitudinal studies have shown that *E. coli* lineages adopt diverse colonization strategies, with both residence-colonization trade-off and niche differentiation processes (19, 20). A better understanding of these adaptive mechanisms, at both genomic and metabolic levels, is essential to disentangle the respective contributions of lifestyle (commensal vs pathogenic), BSI source, and phylogenetic background.

To address this, we analyzed the metabolic diversity of *E. coli* using a well-characterized dataset of commensal and ExPEC strains collected from prospective studies over matched time periods. Our prediction approach leveraged the comprehensive annotation of *E. coli* K-12 pathways, while also extending our capacity to identify poorly characterized or novel pathways through more generalist databases. We confirmed a substantial metabolic diversity across the species, though it is markedly lower than the genomic diversity. We report a pan- and core-metabolism larger and smaller than in previously smaller-scales studies (21, 22), with 487 and 243 pathways identified, respectively. The distribution of pathway frequencies followed a U-shaped curve, opposite to that of gene presence/absence patterns (14, 63), consistent with a faster signal saturation for pathways than for genes. Although this pattern may arise from unidentified pathways, the relatively restricted ecological niche of *E. coli* may involve a limited set of highly specific metabolic functions. Variable pathways were mainly involved in biosynthesis and degradation, in line with previous reports (21, 22), and likely play roles in environmental adaptation and substrate use. Another significant portion of the variable metabolism was associated with O-antigen biosynthesis (22). Of note, the *rfb* gene cluster, involved in the biosynthesis of O-antigen, is a hotspot of recombination in *E. coli* (63) and we previously found major variability in these antigens among the main BSI clones (76), possibly reflecting immune-driven selective pressures.

We then examined metabolic differences associated with strain origin or phylogeny. Consistent with pangenome analysis showing that generalist phylogroups A and B1 harbor larger pangenomes despite smaller genome sizes (23), our results show increased pathway and reaction numbers in phylogroup A. Conversely, strains from urinary tract-related BSI had fewer reactions and pathways. Yet, MCA and heatmap analyses revealed only weak clustering by origin, underscoring the dominant influence of phylogeny. Indeed, more than 61% of urosepsis strains belonged to phylogroup B2, and more than 80% to just 10 STc. Despite the apparent specialization of these clones, we found no specific pathways associated with any BSI portal of entry.

Overall, we observed multiple pathways enriched or depleted in specific phylogroups, nearly all related to degradation, hinting at a role in niche specialization (4, 20). Pathway distribution aligned with phylogroup divisions reported in pangenome analyses (14), suggesting that metabolism in *E. coli* has followed the species evolutionary history. In agreement with earlier studies (21, 24, 25, 63), phylogroup B2 exhibited distinct metabolic features, including frequent absence of degradation pathways for 3-(3-hydroxyphenyl)propanoate, fructoselysine, psicoselysine, and putrescine II.

Zooming in, we also found STc-level associations. For STc10 (phylogroup A), we confirmed previous findings of no specific pathways (25), consistent with its high level of diversity (76). However, in B2 and D STcs, key pathways potentially related to niche adaptation emerged. In STc69, three specific pathways were detected, including S-methyl-5’-thioadenosine degradation l and 5’-deoxyadenosine degradation II, recently identified in ExPEC (66, 67). These pathways enable carbon and sulfur salvage from MTA and 5’-deoxyadenosine via a dihydroxyacetone phosphate shunt. Since these substrates are present in urine and blood, they may support ExPEC growth in extraintestinal environments. Although not significantly enriched in ExPEC overall, these pathways were highly prevalent in the pandemic clone STC131 (81.97%) (Table S8).

Among the 13 B2-specific STc pathways, 10 were involved in degradation of plant-derived products: 3-(3-hydroxyphenyl)propionate degradation in STc131 (19, 23, 25, 77); pectin degradation in STc73, a major STc usually devoid of antibiotic resistance (1); sulfoquinovosyl diacylglycerides and sulfoquinovosyl glycerol degradation in STc14, a pandemic clone whose recent emergence mimics STc131 in several respects (78); and D-apiose degradation in both STc131 and STc14, which are two of the most recently emerged pandemic clones (78, 79). These findings echo Freter’s hypothesis and suggest that such pathways may offer ecological advantages to each of these clones.

Among these pathways, the D-apiose degradation is of particular interest. While its presence in *E. coli* was previously proposed by Carter et al. based on sequence similarity (69), its highly specific distribution in STc131 and STc14 had not been reported. The conserved genomic location in both STcs, its presence in *E. fergusonii*, and the absence of highly similar homologs in other genera suggest an ancient origin in *E. coli*. Of note, the pathway is prevalent in phytopathogenic or plant-associated bacteria, which aligns with the origin of apiose, a branched-chain pentose and a key component of primary cell walls in higher plants (80). Interestingly, several human gut anaerobes, including *Bacteroides* and *Clostridia*, have also been shown to degrade apiose (69), suggesting that it may serve as a relevant nutrient source within the gut niche. Though the pathway lacks synteny with the reference *Pectobacterium* species due to the absence of the D-apiose isomerase, we experimentally confirmed its function via growth assays. Further biochemical studies are warranted to fully characterize the additional enzymes and intermediate compounds in *E. coli*.

Our study has several limitations. First, despite nearly 1,500 genomes, our dataset does not encompass the full diversity of *E. coli*. For example, InPEC and *Shigella* strains were not included. The latter in particular shows drastic metabolic reduction (21, 81). Second, pathway prediction from genome data can misrepresent metabolic capacities due to over- or under-prediction. Nonetheless, our findings are largely consistent with previous studies, and we functionally validated a novel pathway. Third, we could not assess transcriptional or regulatory variation, which may obscure functional metabolic differences. These mechanisms, including substrate affinity, are likely important for niche specialization (5). Last, our binary presence/absence analysis does not consider allelic variation, which could affect enzyme efficiency or specificity. Such analyses require extensive biochemical data not currently available at this scale.

In conclusion, through a large-scale analysis of *E. coli* metabolic pathways, we identified lineage-specific metabolic capacities. While no pathways were strictly associated with pathogenicity or BSI portal of entry, several appeared linked to successful ExPEC clones. These clone-specific metabolic traits may enhance the fitness of major BSI STcs in both gut colonization and extraintestinal infection, allowing them to outcompete resident microbiota and potentially contributing to their emergence and global dissemination. Our findings underscore the strong phylogenetic imprint on *E. coli* metabolism and highlight specific pathways with potential ecological and clinical relevance.

## Acknowledgements

This work was partially funded by the Agence Nationale de la Recherche (ANR): grant DREAM (ANR-20-PAMR-0002). GR was partially supported by a “Poste d’accueil” funded by the “Assistance Publique-Hôpitaux de Paris” (AP-HP) and the “Commissariat a I’energie atomique et aux énergies alternatives” (CEA). We are grateful to Anne Zaparucha for discussion about biochemical aspects of the D-apiose pathway.

