## Supplementary figures for "Identification of specific metabolic capacities associated with major extraintestinal pathogenic *Escherichia coli* lineages"

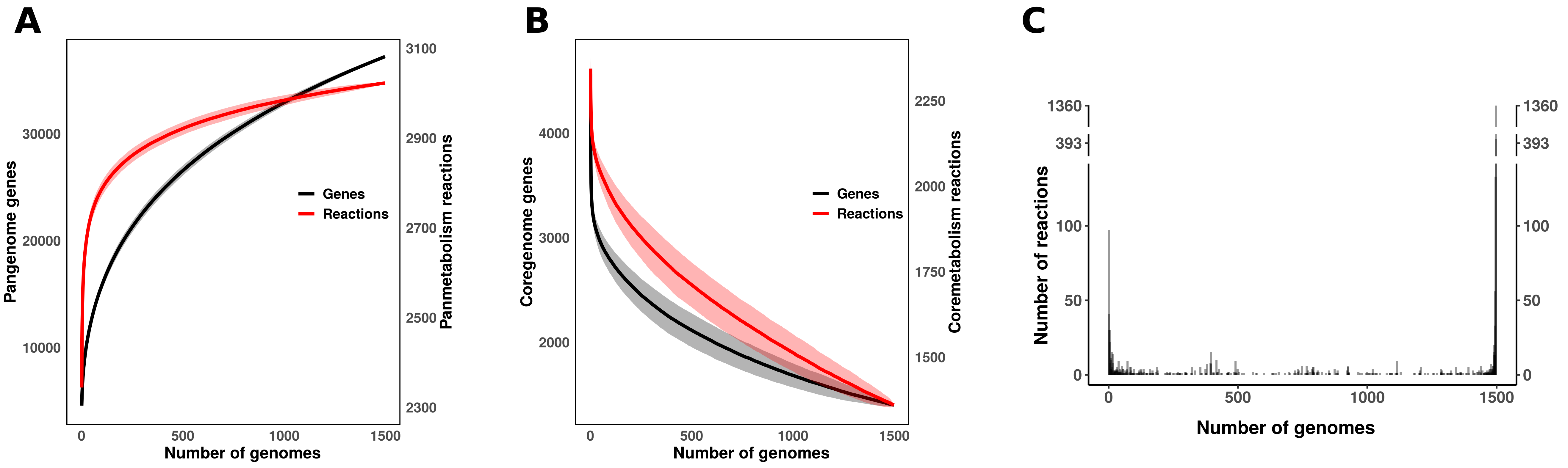


**Figure S1. *E. coli* core- and pan-reactome.** Evolution of (A) the pangenome and panreactome, and (B) the coregenome and corereactome as a function of the number of included genomes. To account for genome variability, 1,000 random permutations were performed at each step of genome addition. The resulting mean number of genes and pathways are shown in black and blue, respectively. Shaded areas represent the standard deviation. (C) Frequency of reactions across the 1,498 genomes analyzed. Reactions on the left side of the graph are present in only one genome (n=97, 3.21% of the panreactome), while those on the right side are found in all genomes (n=1,360, 44.99% of the panreactome). For the sake of readability, the y-axis is broken.

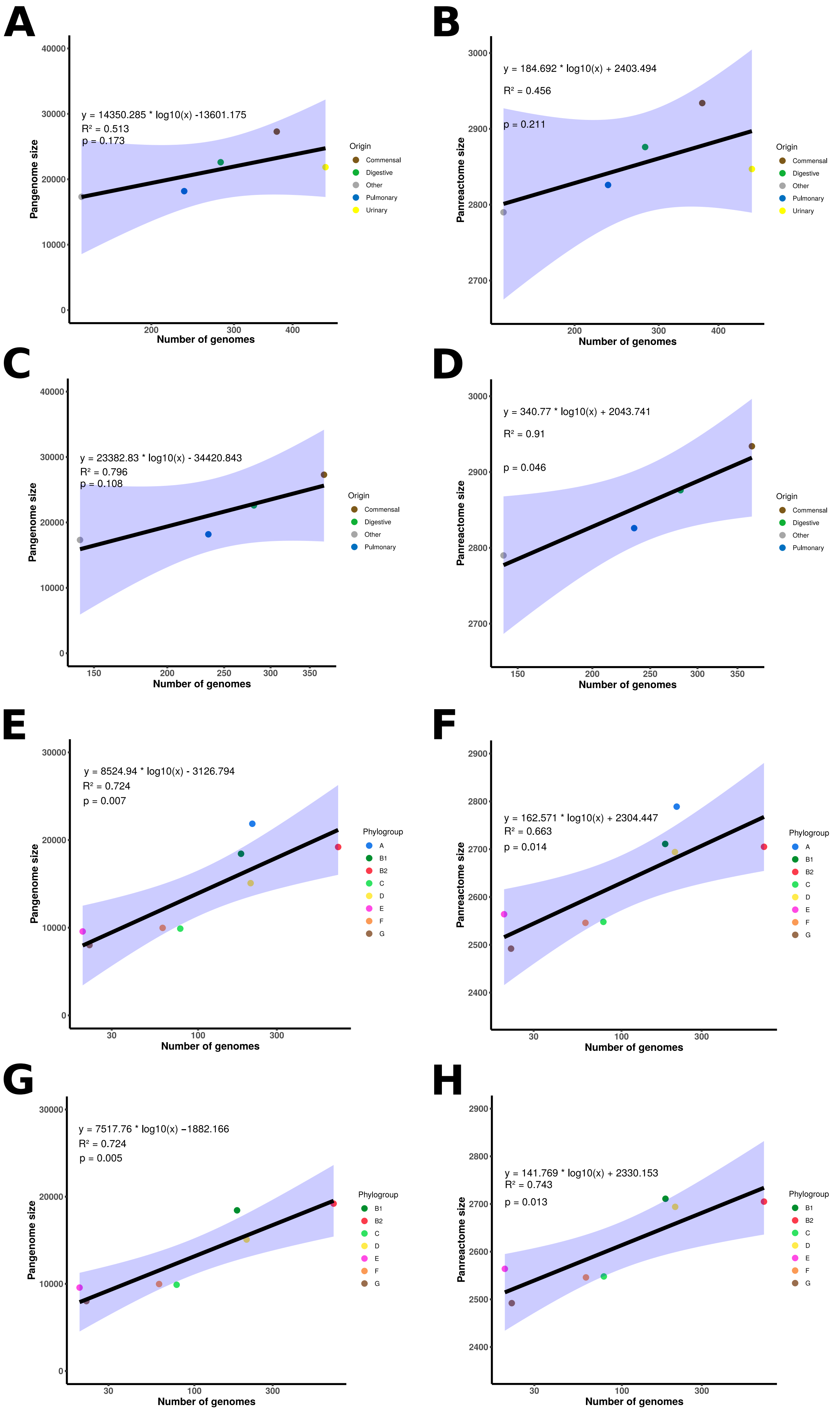


**Figure S2. Correlation between the number of gene families or reactions and genomes.** Total number of gene families and reactions as a function of the number of genomes: (A) and (B) from all sources, (C) and (D) excluding those of urinary origin, (E) and (F) from all phylogroups, and (G) and (H) excluding phylogroup A. Data points are colored according to their origin or phylogroup. The regression line is shown in black, with the 95 % confidence interval in blue. Each graph includes the linear regression equation, the coefficient of determination (R2) and the p-value.

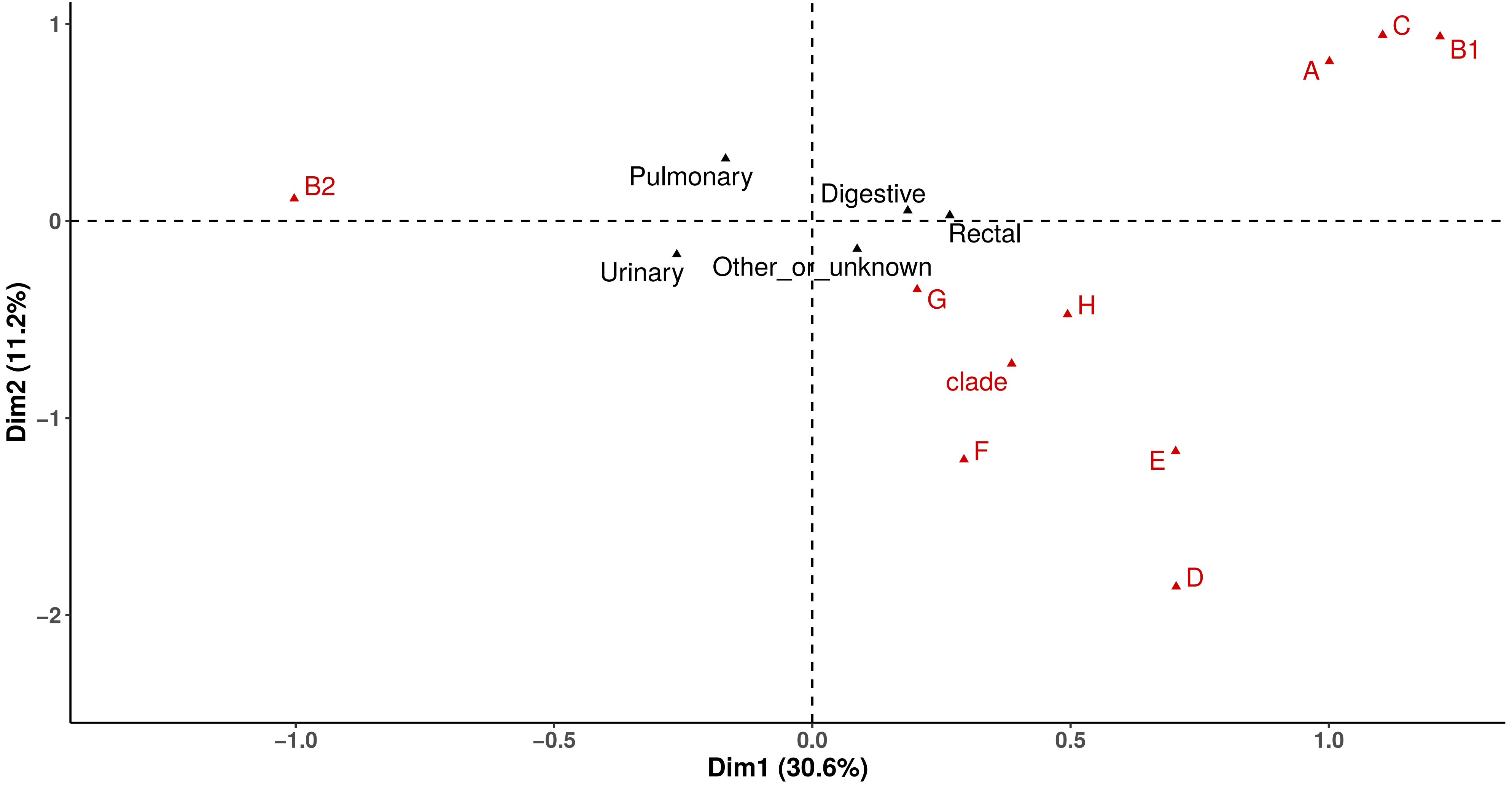


**Figure S3. Multiple correspondence analysis of reaction occurrences among the 1,498 genomes analysed**. The x- and y-axes represent the first dimensions, which together account for 41.8% of the variability. Phylogroups (in red) and origin (in black) are shown as illustrative variables.

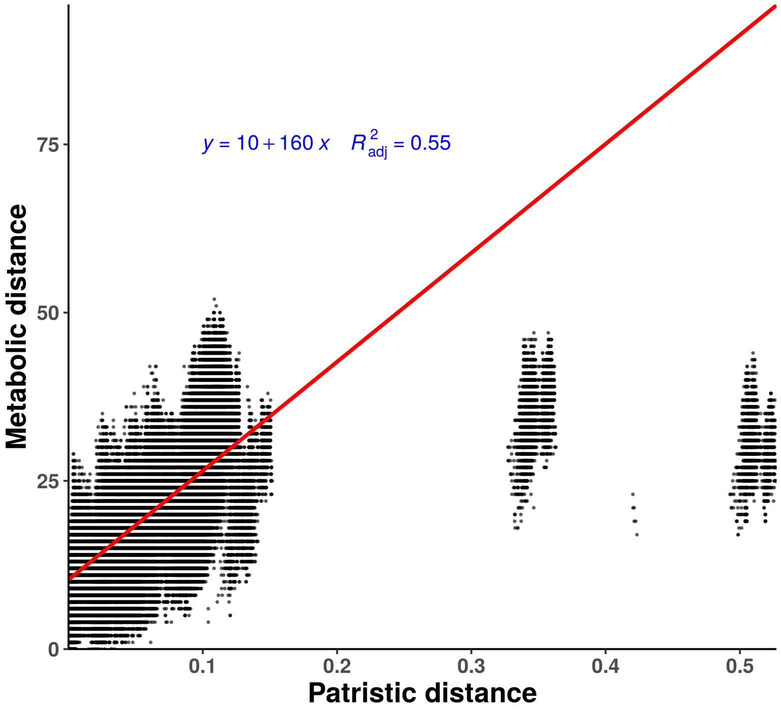


**Figure S4. Patristic distances as a function of metabolic distances between all genome pairs**. Patristic distances represent the branch lengths between genome pairs in the coregene-based phylogenetic tree. Metabolic distances are Manhattan distances computed from binary pathway presence/ absence profiles. Each point represents a pair of genomes. The regression line is shown in red, while the linear regression equation, coefficient of determination (R2) and p-value are shown in blue. The most divergent points on the right side of the plot correspond to comparison between *E. coli* sensu stricto and *Escherichia* clades.

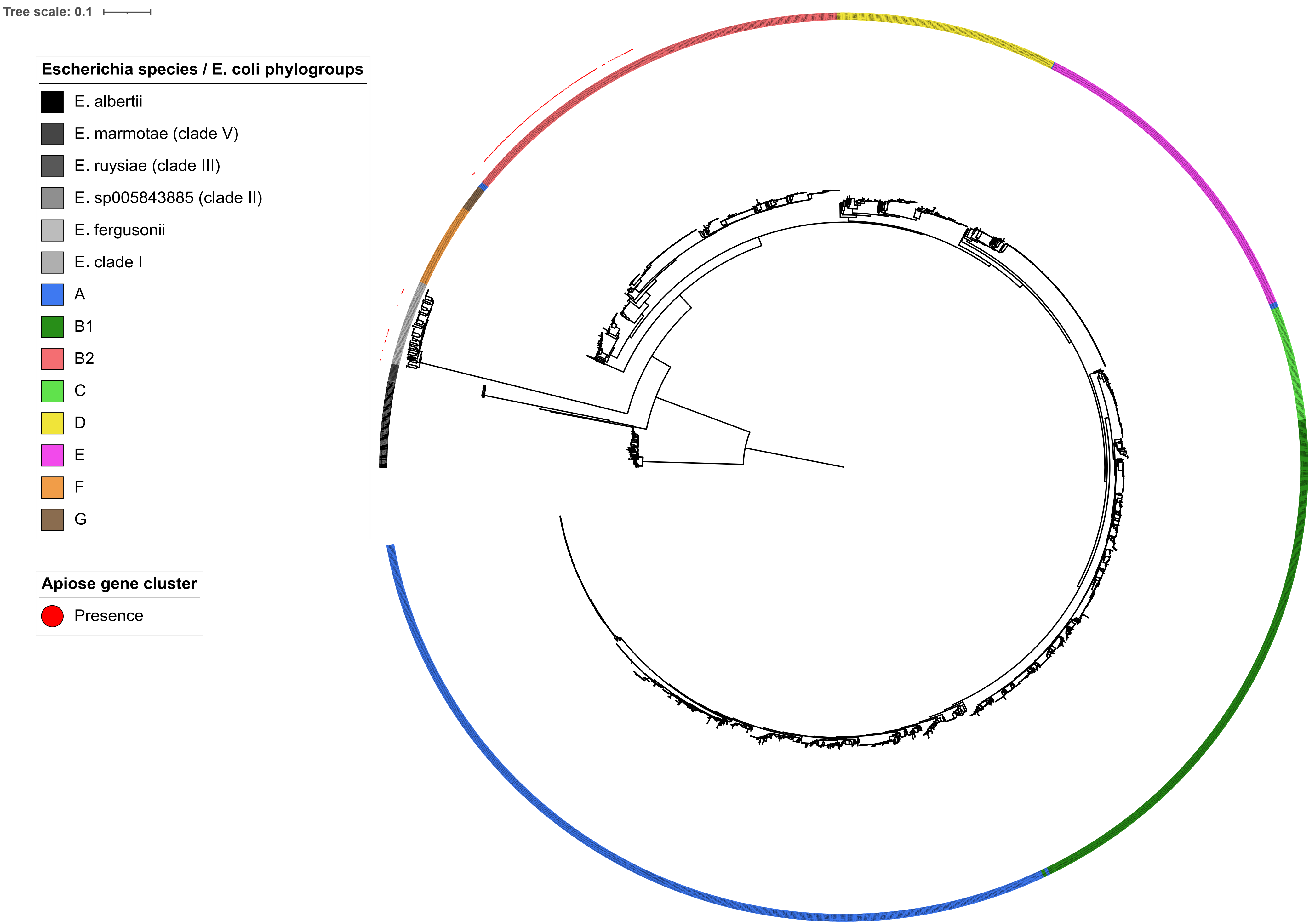


**Figure S5. Maximum likelihood phylogenetic tree based on core gene alignment of complete *Escherichia* genomes from RefSeq**. The tree is rooted on *E. albertii*. The eight main *E. coli* phylogroups are indicated in distinct colors, while *Escherichia* clades and non-coli species are shaded in grey. Red dots indicate the presence of the D-apiose degradation gene cluster. The tree scale indicates the nucleotide substitution per site.

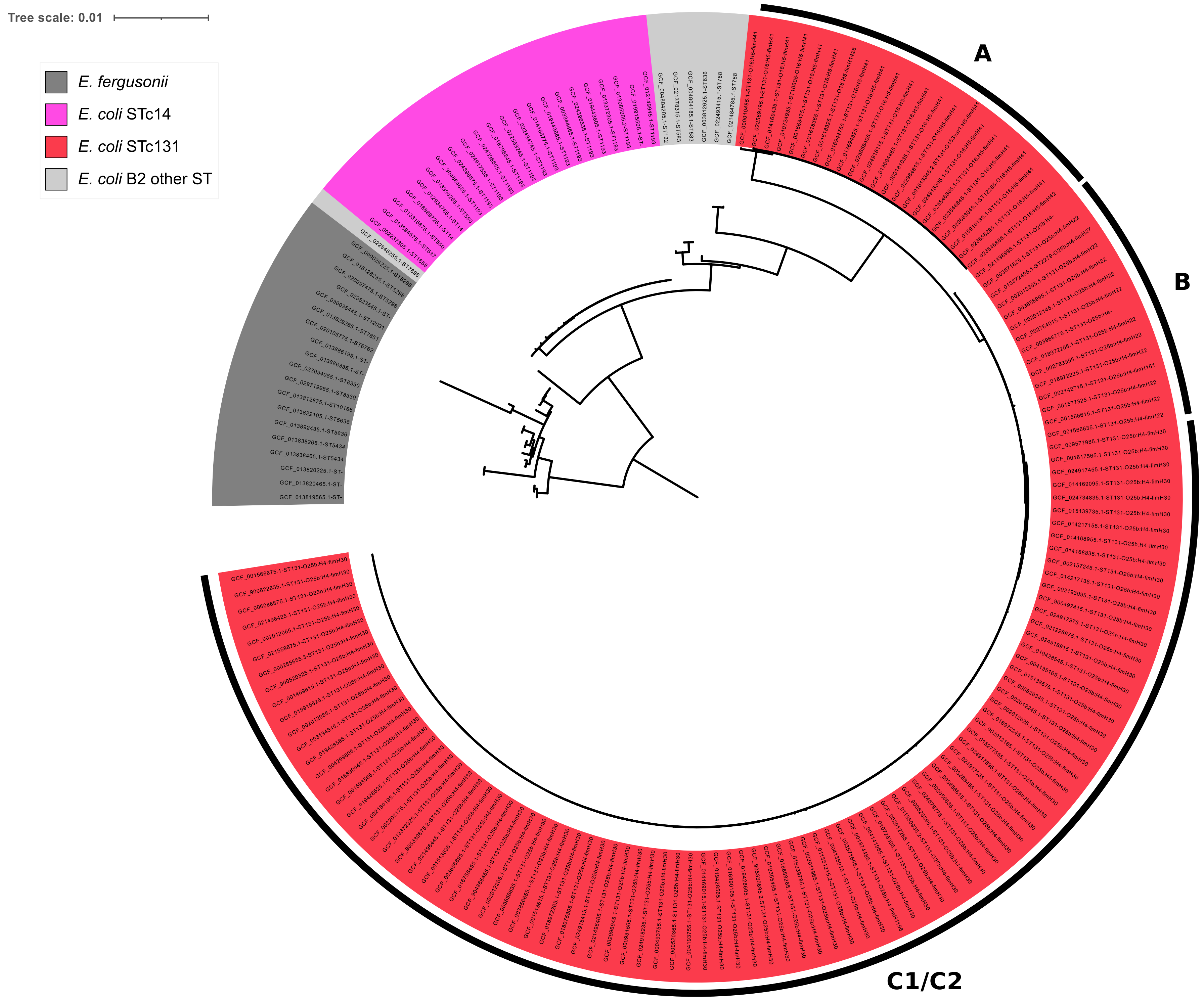


**Figure S6. Maximum likelihood phylogenetic tree computed from the D-apiose degradation gene cluster alignment of complete *Escherichia* genomes from RefSeq.** The tree is rooted on *E. fergusonii*. STc14 and STc131 genomes are highlighted in magenta and red, respectively. Other *E. coli* STs and *E. fergusonii* are shown in light and dark grey, respectively. The outermost ring and corresponding letters denote the clades A, B and C1/C2 of STc131. Leaf labels include the genome name, ST, and O:H-fimH combination for the STc131 genomes. The tree scale indicates the nucleotide substitution per site.

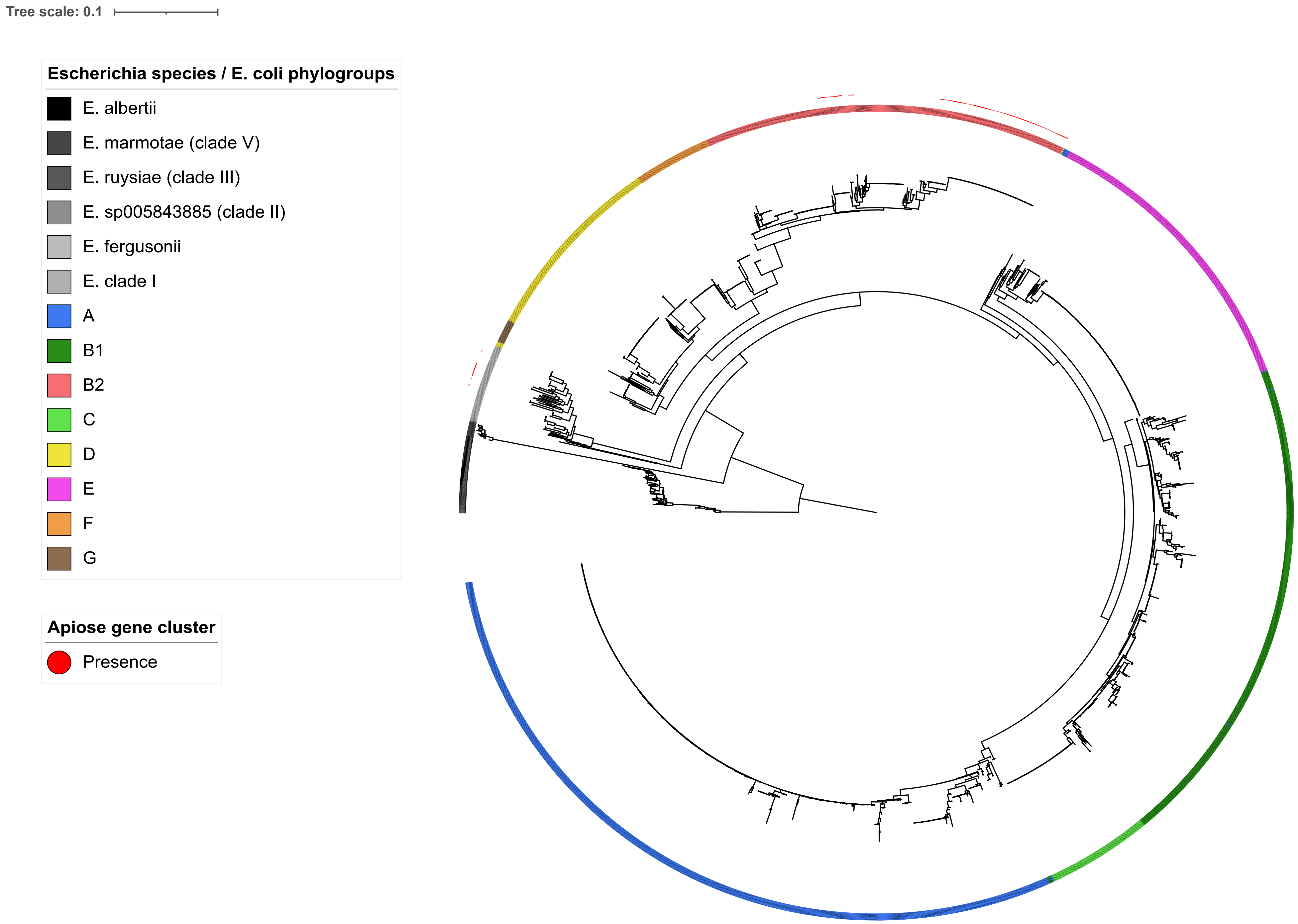


**Figure S7. Maximum likelihood phylogenetic tree computed from the region spanning from *entH* to *cusS* among complete *Escherichia* genomes from RefSeq.** The tree is rooted on *E. albertii*. The eight main *E. coli* phylogroups are indicated in distinct colors, while *Escherichia* clades and non-coli species are shaded in grey. Red dots indicate the presence of the D-apiose degradation gene cluster. The tree scale indicates the nucleotide substitution per site.
